# *Drosophila melanogaster* Set8 and L(3)mbt function in gene expression independently of histone H4 lysine 20 methylation

**DOI:** 10.1101/2024.03.12.584710

**Authors:** Aaron T. Crain, Megan B. Butler, Christina A. Hill, Mai Huynh, Robert K. McGinty, Robert J. Duronio

## Abstract

Mono-methylation of Lysine 20 of histone H4 (H4K20me1) is catalyzed by Set8 and thought to play important roles in many aspects of genome function that are mediated by H4K20me-binding proteins. We interrogated this model in a developing animal by comparing in parallel the transcriptomes of *Set8^null^*, *H4^K20R/A^*, and *l(3)mbt* mutant *Drosophila melanogaster*. We found that the gene expression profiles of *H4^K20A^* and *H4^K20R^* larvae are markedly different than *Set8^null^* larvae despite similar reductions in H4K20me1. *Set8^null^* mutant cells have a severely disrupted transcriptome and fail to proliferate *in vivo*, but these phenotypes are not recapitulated by mutation of *H4^K20^* indicating that the developmental defects of Set8*^null^* animals are largely due to H4K20me1-independent effects on gene expression. Further, the H4K20me1 binding protein L(3)mbt is recruited to the transcription start sites of most genes independently of H4K20me even though genes bound by L(3)mbt have high levels of H4K20me1. Moreover, both Set8 and L(3)mbt bind to purified H4K20R nucleosomes in vitro. We conclude that gene expression changes in *Set8^null^* and *H4^K20^* mutants cannot be explained by loss of H4K20me1 or L(3)mbt binding to chromatin, and therefore that H4K20me1 does not play a large role in gene expression.

## Introduction

The prevailing model for how histone post-translational modifications (PTMs) regulate gene expression is that “writer” enzymes establish where and when histone PTMs are deposited in the genome and “reader” proteins bind to these PTMs to activate or repress transcription by modulating the recruitment of transcription factors (Jenuwein and Allis 2001; Kouzarides 2007; Rothbart and Strahl 2014; Strahl and Allis 2000). Support for this model primarily comes from experiments that infer histone PTM function through genetic manipulation of individual writers or readers (Henikoff and Shilatifard 2011; Kouzarides 2007). However, this approach has a major caveat: most writers have non-histone substrates and readers interact with many proteins other than modified histones (Clarke 2013; Zhang et al. 2015). Determining the roles of specific histone PTMs in genome function and development would benefit from a comparative analysis of mutations in a writer, its target histone residue, and a reader of the modified histone residue in a single animal model system. Such a combined approach is only feasible in *Drosophila melanogaster* where the generation of histone mutant genotypes can be achieved (McKay et al. 2015). To better understand the role of H4K20 methylation in genome function, we used genomic approaches in *Drosophila* to examine in parallel gene expression phenotypes resulting from mutations of histone H4 lysine 20 (H4K20), the H4K20-specific mono-methyltransferase Set8, and an H4K20me binding protein, L(3)mbt.

Methylation of histone H4 lysine 20 (H4K20me) has been implicated in the regulation of DNA replication (Li et al. 2016; Hayashi-Takanaka et al. 2021; Shoaib et al. 2018; Brustel et al. 2017; Pellegrino et al. 2017; Huen et al. 2008; Jørgensen et al. 2007; Tardat et al. 2007; Takawa et al. 2012), gene expression (Kalakonda et al. 2008; Karachentsev et al. 2005; Lv et al. 2016; Morgan and Shilatifard 2020; Veloso et al. 2014; Yang et al. 2012; Yu et al. 2019; Barski et al. 2007; Beck et al. 2012; Congdon et al. 2010), and DNA damage repair (Sakaguchi and Steward 2007; Dulev et al. 2014; Jørgensen et al. 2007) in both humans and flies, suggesting an evolutionarily conserved role for H4K20me in critical aspects of genome function and stability. Indeed, the human H4K20 mono-methyltransferase Set8 (aka KMT5A) can replace essentially all developmental functions of *Drosophila* Set8 (Crain et al. 2022). Different methylation states of H4K20 (i.e. K20me1, K20me2, and K20me3) are thought to mediate different genomic functions. Set8 catalyzes H4K20me1, the preferred substrate for subsequent di- and tri- methylation by Suv4-20 enzymes (Weirich et al. 2016; Yang et al. 2008; Southall et al. 2014; Schotta et al. 2008). Thus, loss of Set8 depletes all H4K20 methyl states and results in pleiotropic phenotypes and lethality in both flies and mice (Karachentsev et al. 2005; Oda et al. 2009; Crain et al. 2022). Therefore, current models in the field posit that Set8 regulates genome functions mainly through deposition of H4K20me1.

The genomic functions of H4K20me are thought to be mediated by proteins that recognize various methylation states of H4K20, including lethal (3) malignant brain tumor (L(3)mbt), Orc1, 53BP1, enhancer of zeste (E(z)), and the Msl complex (Kim et al. 2010; Kuo et al. 2012; Sakaguchi et al. 2012; Tuzon et al. 2014; Weaver et al. 2019; Kalakonda et al. 2008). Loss of L(3)mbt in *Drosophila* results in a somatic to germline shift in gene expression, leading to over-proliferation of brain tissue (Bonasio et al. 2010; Janic et al. 2010). L(3)mbt contains three MBT domains that share homology with the “Royal” family of chromatin interacting proteins and bind H4K20me1/2 *in vitro* (Kalakonda et al. 2008; Qi et al. 2010; Sakaguchi et al. 2012; Trojer et al. 2007; Min et al. 2007; Maurer-Stroh et al. 2003). L(3)mbt interacts *in vivo* with several partner proteins as a member of the LINT and MybMuvB/dREAM complexes, executing transcriptional repression seemingly independently of H4K20me (Blanchard et al. 2014; Yamamoto-Matsuda et al. 2022; Coux et al. 2018; Meier et al. 2012). Thus, the connection between L(3)mbt histone binding of H4K20me and its regulation of gene expression is incompletely understood. Numerous previous studies suggest functional connections between H4K20me, Set8, and L(3)mbt. For instance, depletion of Set8 or L(3)mbt results in a loss of H4K20me1 and causes defects in DNA replication, chromatin organization, and transcription, implying that H4K20me1 plays a causal role in these processes (Beck et al. 2012; Sakaguchi et al. 2012). However, Set8 has non-histone substrates [e.g. p53 (Shi et al. 2007) and PCNA (Takawa et al. 2012) and non-catalytic functions [(e.g. in cell cycle entry, (Yin et al. 2008; Zouaz et al. 2018), and L(3)mbt co-localizes promiscuously with several different mono- and di- methylated histone residues (Blanchard et al. 2014). Furthermore, *Drosophila* mutants expressing unmodifiable H4K20A or H4K20R are phenotypically distinct from *Set8^null^* mutants (Crain et al. 2022), and loss of L(3)mbt function does not recapitulate many of the cell proliferation defects in *Set8^null^* animals, despite a 60% reduction in H4K20me1 (Sakaguchi et al. 2012). Thus, whether and how H4K20me mediates the functions of Set8 and L(3)mbt remains unclear.

H4K20me1’s role in gene expression in mammals and flies is also not easily reconciled into a simple model. In mammalian cells, H4K20me1 is found in the body of actively transcribed genes (Barski et al. 2007; Beck et al. 2012), suggesting a role for H4K20me1 in stimulating transcription (Kapoor-Vazirani and Vertino 2014; Nikolaou et al. 2017; Veloso et al. 2014; Shoaib et al. 2021) that was also suggested for *Drosophila* H4K20me1 (Huang et al. 2021; Yu et al. 2019; Lv et al. 2016). Conversely, H4K20me1 has been implicated in chromatin compaction (Lu et al. 2008) and gene repression (Congdon et al. 2010; Abbas et al. 2010; Liu et al. 2010; Kalakonda et al. 2008), either via its association with the transcriptional repressor L(3)mbt or its localization in inactive regions of the genome (Kalakonda et al. 2008; Nishioka et al. 2002; Sakaguchi et al. 2012). Here we determined whether the gene expression phenotypes arising upon removal of Set8 or L(3)mbt can be attributed to loss of H4K20me1. By leveraging null and hypomorphic alleles of *Set8* and *l(3)mbt* with our ability to engineer fully histone mutant genotypes (e.g. *H4^K20A^* and *H4^K20R^*) we demonstrate that the major roles of Set8 and L(3)mbt in gene expression and cell proliferation do not require H4K20me. Our data suggest that phenotypes resulting from mutating H4K20 are due to effects on H4 binding proteins rather than loss of H4K20me.

## Results

### H4K20me1 is correlated with actively transcribed genes in *Drosophila* larva

Although H4K20me1 ChIP-seq datasets exist for *Drosophila* (Lv et al. 2016; modENCODE Consortium et al. 2010), the genome-wide relationship between H4K20me1 and gene expression has not been elucidated in flies. To determine this relationship, we first performed CUT&RUN genomic occupancy profiling in wild type Oregon-R third instar wing imaginal discs using an H4K20me1-specific antibody. We measured enrichment of H4K20me1 signal over a no primary antibody control using a sliding window method and found H4K20me1 signal in broad peaks primarily enriched in gene bodies, consistent with previous studies in mammalian cells (**Figure 1A, B)** (Barski et al. 2007; Beck et al. 2012). Using a metagene analysis we found that H4K20me1 signal overlapped by at least one bp with 5366 protein-coding genes, and that the amount of overlap varied among genes. For instance, 2309 protein-coding genes were covered at least 50% by H4K20me1 and 1439 had 75% coverage (the H4K20me1 “HIGH” category) (**Figure 1C, D**). The remaining 11,707 protein-coding genes did not have H4K20me1 signal that was statistically enriched compared to control or were covered less than 50% by an H4K20me1 peak (the H4K20me1 “LOW” category) (**Figure 1C, D**). We also found that the proportion of H4K20me1 coverage of each gene correlated positively with the amount H4K20me1 signal at each gene (**Figure 1D**).

**Figure 1.**
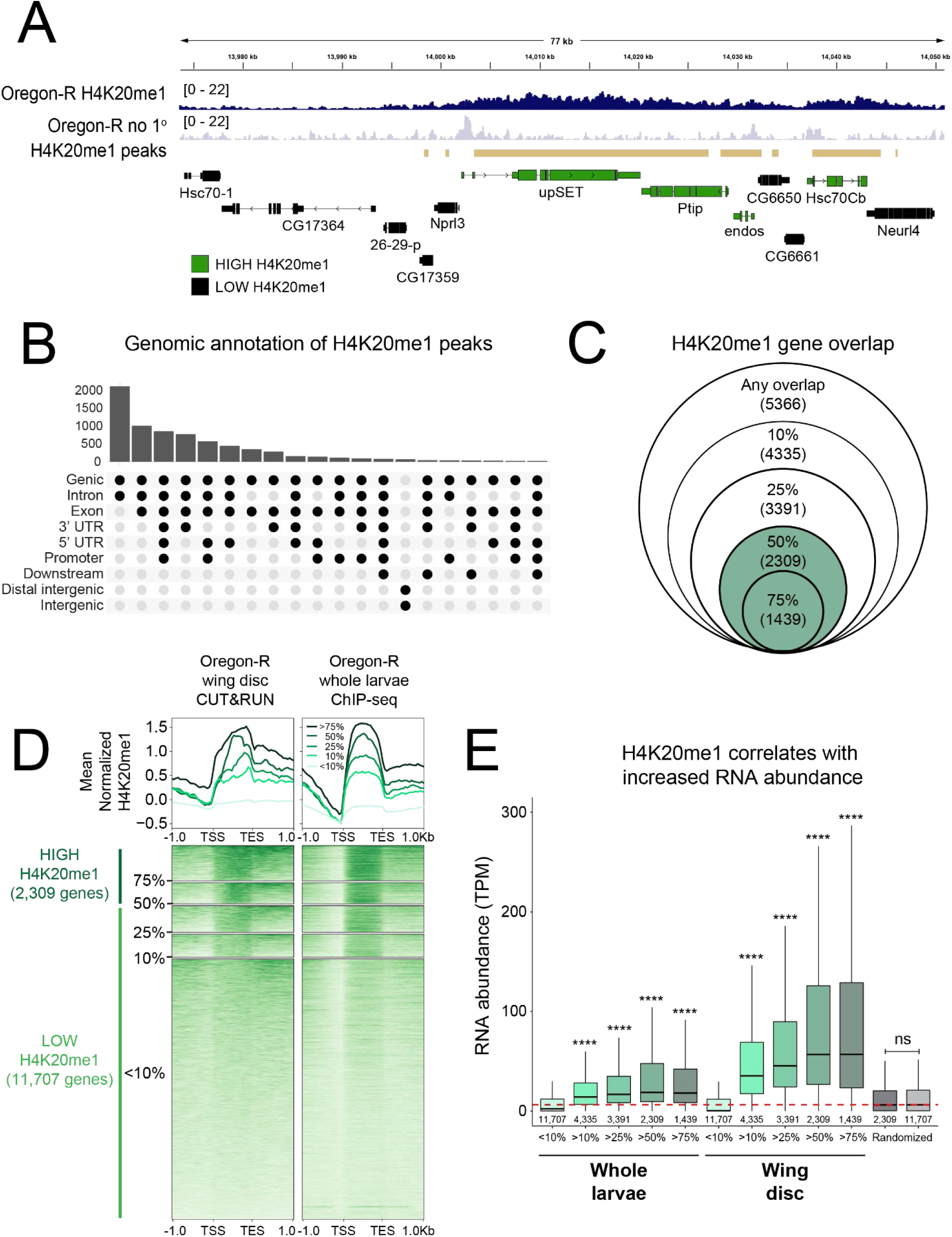
H4K20me1 is enriched in gene bodies in *Drosophila melanogaster.* **A)** Genome browser shot of a representative locus depicting H4K20me1 CUT&RUN signal. Gold bars indicate peaks of H4K20me1 signal in wild type Oregon-R relative to no primary antibody control as defined by merged 150bp sliding windows. **B)** Upset plot depicting frequency of H4K20me1 peaks overlapping specific genomic features. Histogram (top) indicates the number of peaks that overlap each of the genomic features shown below each bar. **C)** Venn diagram depicting the number of genes covered by H4K20me1 peaks. Outer circle indicates >1bp overlap of a gene covered by an H4K20me1 peak, followed by 10%, 25%, 50%, and 75% coverage. Green indicates our H4K20me1 “HIGH” category. **D)** Heatmap (bottom) and summary (top) metaplot of H4K20me1 coverage in gene bodies scaled to 1kb as well as plus and minus 1kb unscaled sequence. The plots are organized by gene sets defined in Figure 1C for Oregon-R wing disc CUT&RUN in this study (left) and Oregon-R whole larvae ChIP-seq from the modEncode database (right). Genes with >50% overlap with H4K20me1 peaks are considered HIGH H4K20me1 (Dark green, left side). Genes with <50% overlap with H4K20me1 are considered LOW H4K20me1 (Light green, left side). Each gene row is ordered by mean H4K20me1 in the wing disc dataset. Summary plot depicts mean signal at each position in the metaplot (50bp bins) for each set of genes. **E)** Boxplot of average RNA abundance (transcripts per million, TPM) of genes in each of the gene sets defined in Figure 1C in whole larvae RNA-seq (this study), wing disc RNA-seq (Armstrong et al. 2018), or a random set of genes of indicated size. Red dotted line indicates median RNA abundance in whole larvae. Significance determined by Wilcoxon Sum Rank Test with Benjamin-Hochberg multiple testing correction. **** indicates p < 0.0001 for each indicated set compared to both genes covered by <10% H4K20me1 peak and the randomized gene set.

We then performed RNA-sequencing of Oregon-R whole third instar larvae to assess whether H4K20me1 correlates with RNA abundance. Whole animal RNA-seq was required due to the difficulty of obtaining enough material from individual wing discs of some mutant genotypes (see below). Our H4K20me1 CUT&RUN data in wing disc tissue correlates well with a previous modENCODE H4K20me1 ChIP-seq dataset in whole larvae, suggesting the distribution of H4K20me1 within genes is consistent across tissues (**Figure 1D**). We found that genes in the H4K20me1 HIGH category were more highly expressed compared to genes in the H4K20me1 LOW category in both whole larval RNA-seq (this study) and wing disc RNA-seq data (Armstrong et al. 2018) (**Figure 1E**). HIGH H4K20me1 genes and LOW H4K20me1 genes were also significantly different than random gene sets of equal size (**Figure 1E**). Together, these data suggest that H4K20me1 deposition in gene bodies is positively correlated with active transcription in developing *Drosophila* larvae.

### Modification of H4K20 is not required for transcription

Since H4K20me1 is positively associated with gene expression, we asked whether loss of H4K20me1 would result in downregulation of genes with H4K20me1 by performing H4K20me1-specific CUT&RUN in *Set8^null^*third instar wing imaginal discs. Consistent with previous studies, *Set8^null^*mutants have a reduction in H4K20me1 levels across gene bodies throughout the genome (**Figure 2A, B**) (Beck et al. 2012; Karachentsev et al. 2005; Nishioka et al. 2002). Surprisingly, *Set8^null^*wing discs from wandering third instar larvae have residual H4K20me1 as assessed by CUT&RUN (**Figure 2A**) despite *Set8^null^* larval brains lacking H4K20me1 signal by western blot (Crain et al. 2022). CUT&RUN is a more sensitive assay and others have reported H4K20me1 in *Set8^null^* third instar wandering salivary glands (Karachentsev et al. 2005). We conclude that previous studies using western blots lacked the sensitivity to detect low H4K20me1 levels in *Set8^null^*mutants.

**Figure 2.**
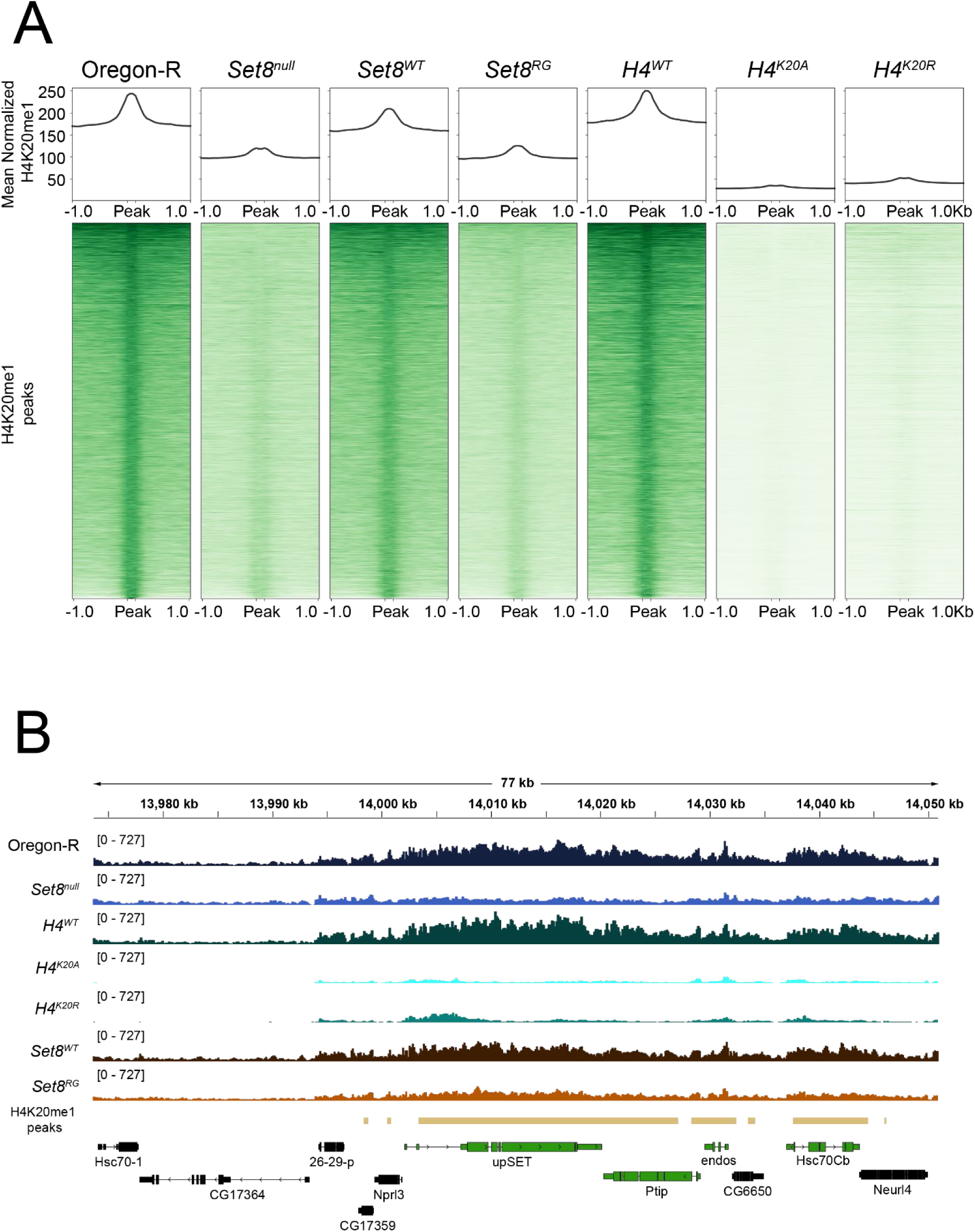
Loss of H4K20me in *Set8* and *H4^K20^* mutants. **A)** Heat (bottom) and summary (top) metaplots of spike-in normalized H4K20me1 signal at H4K20me1 peaks in indicated genotypes. Each peak is scaled to 200bp and flanked by 1kb of unscaled sequence. The summary plot depicts mean signal at each position in the metaplot (50kb bins) for all peaks. **B)** Representative locus depicting H4K20me1 CUT&RUN coverage in wing discs. Gold bars indicate peaks of H4K20me1 signal in Oregon-R relative to no primary antibody control as defined by merged 150bp sliding windows. Green and black genes indicate H4K20me1 “HIGH” and “LOW” categories, respectively.

To ask whether the decreased H4K20me1 levels in *Set8^null^*mutants were associated with a change in gene expression, we performed total RNA-seq in *Set8^null^* larvae and compared these data to the Oregon-R whole larvae RNA-seq dataset described in Figure 1. We chose to assess gene expression in whole mutant larvae even though we measured H4K20me1 in wing discs because *Set8^null^* animals have small, morphologically perturbed wing discs containing few cells, (Karachentsev et al. 2005) making it challenging to obtain enough high-quality material for RNA-seq even though we obtained enough for CUT&RUN. Moreover, there is no evidence for tissue-specific patterns of H4K20me1 deposition in chromatin (see **Figure 1**). We therefore considered the loss of H4K20me1 we observed in wing discs to be representative of other tissues.

Differential expression analysis of *Set8^null^* relative to Oregon-R wild type control revealed that *Set8^null^* animals had 1,491 differentially expressed genes (DEGs, 962 up-regulated, 529 down-regulated), indicating that loss of Set8 significantly disrupts the *Drosophila* transcriptional program (**Figure 3A, Supplemental Table 1**). The majority of DEGs (962, 65% of total DEGs) are upregulated, and only 86 *Set8^null^* DEGs (18 up-regulated and 68 down-regulated) are in the HIGH H4K20me1 category (**Figure 3A**, dark dots). These 86 represent 5.8% of all *Set8^null^* DEGs and only 3.7% of genes in the HIGH H4K20me1 category, indicating that loss of H4K20me1 is not predictive of either an increased or decreased change in gene expression (**Figure 3A**, **Supplemental Table 2**). Similarly, only two genes in the H4K20me1 HIGH category are down-regulated in animals expressing catalytic-deficient Set8 [*Set8^R634G^*, hereafter *Set8^RG^*, (Crain et al. 2022) (**Figure 3D**), despite a similar reduction in H4K20me1 levels by CUT&RUN (**Figure 2A, B**). These data indicate that H4K20me1 does not play a causal, global role in transcriptional control.

**Figure 3.**
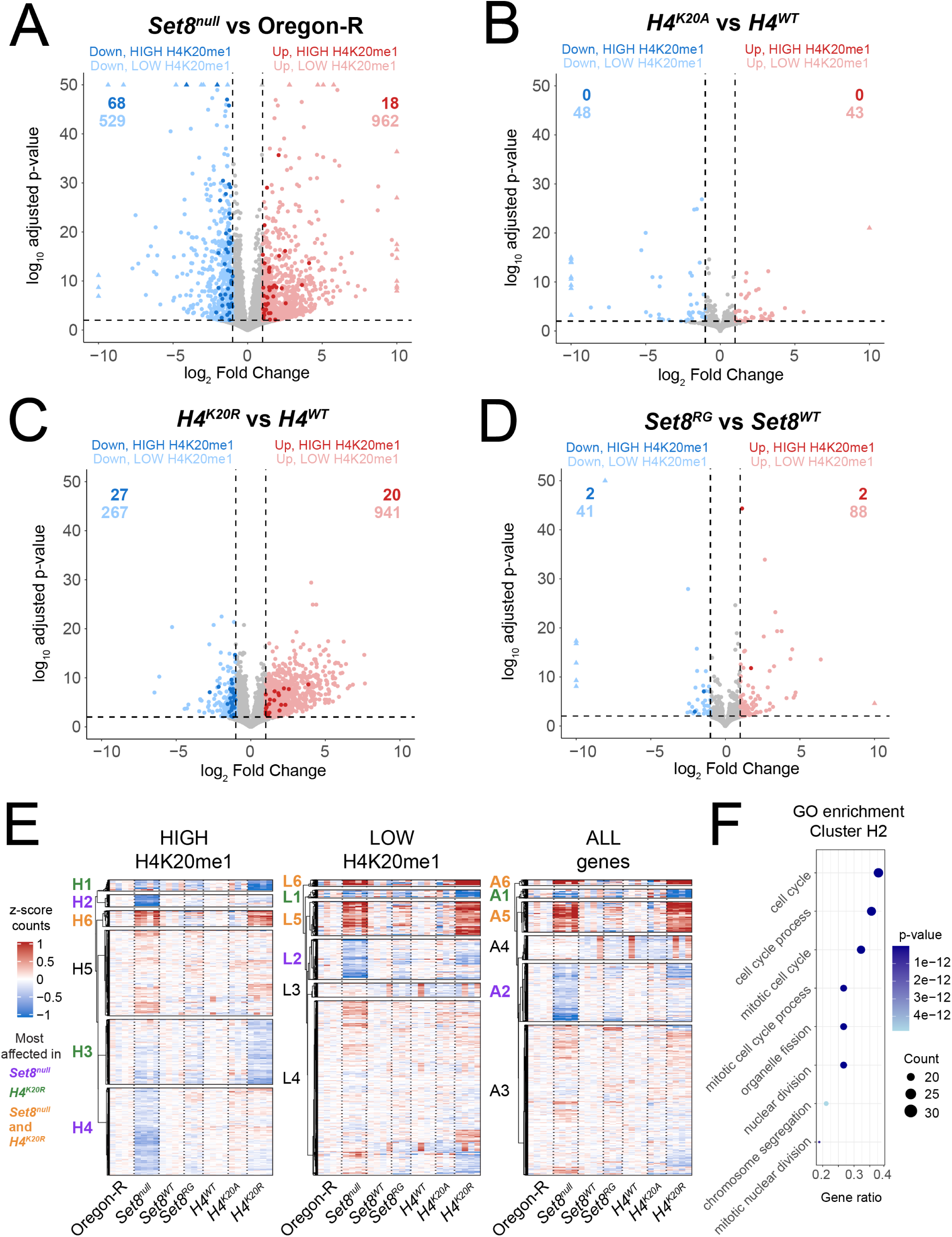
The *Set8* and *H4^K20^* mutant transcriptomes differ. **A-D)** Volcano plots depicting the relationship between log_2_FoldChange (x-axis) and log_10_ adjusted p-value (y-axis) of gene expression in indicated comparisons. Blue dots indicate significantly downregulated genes (log_2_FC < −1, FDR < 0.01) and red dots indicate significantly upregulated genes (log_2_FC > 1, FDR < 0.01). Darker shaded dots indicate genes in the HIGH H4K20me1 category. **E)** Clustered heatmaps of average centered normalized counts. Cluster identifiers are colored based on whether expression of genes within that cluster are most affected in *Set8^null^* (purple), *H4^K20R^* (green), or both *Set8^null^* and *H4^K20R^* (orange). Black cluster identifiers indicate no enrichment of gene expression changes in any genotype. **F)** GO enrichment summary for cluster H2. Top eight GO terms are shown on the Y-axis indicates. The x-axis indicates ratio of genes from cluster H2 that intersect with each GO term over all genes in cluster H2. Size of the dot indicates number of intersecting genes and color indicates p-value of each term. Full list of GO terms for clusters in Figure 3G can be found in **Supplemental Tables 1.1-1.3**.

To investigate the role of H4K20me1 in gene expression more directly, we performed H4K20me1- specific CUT&RUN in wing imaginal discs and RNA-sequencing of third instar whole larvae in *H4K20* mutant (*H4^K20A^* and *H4^K20R^*) and *H4^WT^* control genotypes. Both *H4^K20A^* and *H4^K20R^* have a strong reduction of H4K20me1 genome wide, despite the presence of replication-independent *His4r* in these genotypes (**Figure 2A**). *His4r* is a single copy gene located outside of the replication-dependent histone gene array, and its expression is not replication coupled but encodes a protein with an amino acid sequence identical to replication-dependent H4. We and others have shown previously that *His4r* can partially compensate for the absence of replication-dependent H4 (Crain et al. 2022; Armstrong et al. 2018; Copur et al. 2018; Faragó et al. 2021). Nevertheless, our H4K20me1 CUT&RUN data indicate that His4r is a minor contributor to H4K20me1 amounts in the genome.

Despite levels of H4K20me1 lower than in *Set8^null^*(**Figure 2A**), *H4^K20A^* animals have only a small number of DEGs (91 total; 43 up-regulated, 48 down-regulated) relative to control (**Figure 3B, Supplemental Table 1**). Remarkably, no *H4^K20A^* DEGs are in the HIGH H4K20me1 category (**Figure 3B**, **Supplemental Table 2**). *H4^K20R^* animals have a larger number of DEGs (1208 total; 941 up-regulated, 267 down-regulated) compared to *H4^K20A^* (**Figure 3C, Supplemental Table 1**). Still, only 3.9% (47) of *H4^K20R^*DEGs are in the HIGH H4K20me1 category, and only 57.4% (27) of those DEGs are down-regulated in *H4^K20R^* (**Supplemental Table 2**). Given that H4K20me1 levels are severely depleted in *H4^K20A^* and *H4^K20R^*animals, these data suggest that transcript levels of most genes are not sensitive to reduction of H4K20me1 but are instead sensitive to the residue identity at position 20 on the H4 tail. Together, these data further emphasize that the level of H4K20me1 is not causal for gene expression, despite H4K20me1 enrichment at transcriptionally active genes (**Figure 1**). They also emphasize that the differences in both the gene expression and developmental phenotypes (Crain et al. 2022) of *H4^K20A^* and *H4^K20R^* mutants arise from something unrelated to H4K20 methylation.

### *H4^K20A/R^* and *Set8^null^* gene expression profiles are distinct

Although H4K20me1 is not required for expression of most genes, a subset of genes enriched for H4K20me1 might depend on H4K20me1 and be drivers of gene expression cascades. We addressed this question by comparing normalized gene counts in *Set8^null^*, *H4^K20A^*, and *H4^K20R^* mutants through k-means clustering. By applying this method first to genes in the HIGH H4K20me1 category we found distinct clusters of down-regulated genes: two that were specific to *Set8^null^* (**Figure 3E**, clusters H2 and H4) and two containing genes that were most affected in *H4^K20R^* (**Figure 3E**, clusters H1 and H3). We also found that most upregulated genes in the HIGH H4K20me1 category were shared between *Set8^null^* and *H4^K20R^*(**Figure 3E**, cluster H6).

As with the HIGH H4K20me1 category, we found that the LOW H4K20me1 category of genes contained clusters of down-regulated genes that were most affected in *Set8^null^* (cluster L2) or *H4^K20R^* (**Figure 3E**, cluster L1). We also found clusters that were upregulated in both *Set8^null^* and *H4^K20R^* but not in *H4^K20A^*or *Set8^RG^* **(Figure 3E**, clusters L5 and L6), suggesting that changes in the expression of genes in these clusters do not arise from loss of H4K20me1. Moreover, even when we considered ALL genes independently of H4K20me status, the resulting cluster patterns were remarkably consistent to when we considered only the HIGH or LOW H4K20me1 categories. Finally, the small number of *H4^K20A^* or *Set8^RG^* up- or down-regulated genes did not show a clustering pattern like either *Set8^null^*or *H4^K20R^* for either the H4K20me1 HIGH or LOW categories. Thus, when comparing transcriptomes using either DEGs or k-means clustering of normalized counts each mutant displays a distinct gene expression profile that cannot easily be explained by loss of H4K20me.

Since we observed distinct gene expression patterns in *Set8^null^* and *H4^K20R^* mutants, we were curious if genes in each of the identified clusters were enriched in specific biological processes. Therefore, we performed gene ontology (GO) enrichment analysis on each DEG cluster and grouped significant GO terms by semantic similarity (**Supplemental Figures 1.1-1.3, Supplemental Tables 1.1-1.3**). We were especially interested in clusters H2 and H4 that contained a subset of down-regulated genes in the HIGH H4K20me1 category that were specific to *Set8^null^*. Cluster H2 is enriched for GO terms related to the cell cycle, cell proliferation, cell differentiation, and development (**Figure 3F**, **Supplemental Figure 1.1, Supplemental Table 1.1**). Cluster H4 is enriched for GO terms related to gene expression, RNA processing/splicing, and histone modification (**Supplemental Figure 1.1, Supplemental Table 1.1**). Clusters H2 and H4 were also represented in clusters of genes with LOW H4K20me1 and all genes (L2 and A2), suggesting that these changes in gene expression are H4K20me1-independent (**Supplemental Figure 1.2-1.3, Supplemental Table 1.2-1.3, Figure 3E**). Clusters H1 and H3, containing genes down-regulated primarily in *H4^K20R^*, are enriched for GO terms related to ubiquitin-dependent catabolism and development (H1) as well as metabolism, cell differentiation, cell signaling, growth, and development (H3) (**Supplemental Figure 1.1, Supplemental Table 1.1**). Although both *Set8^null^*-specific (H2 and H4) and *H4^K20R^*-specific (H1 and H3) clusters are enriched for genes involved in development and cell differentiation, the mis-regulated genes themselves are distinct. Together, we conclude that the *Set8^null^*and *H4^K20^* mutant gene expression profiles are overlapping yet contain unique features, and thus are not primarily driven by loss of H4K20me1.

### Cell proliferation defects upon removal of Set8 are largely independent of loss of H4K20me

One of the unique features of the *Set8^null^* transcriptome is the down-regulation of genes involved in the cell cycle (**Figure 3E, F**, clusters H2, L2, A2). We also found GO terms related to growth in *H4^K20R^* (**Figure 3E**, cluster H3). We therefore assessed the cell proliferation capacity of *Set8^null^* versus *H4^K20^* mutant cells. We generated mosaic tissue in the *Drosophila* eye by inducing mitotic recombination with the FLP/FRT system using *eyeless*-*FLP*, which expresses FLP recombinase throughout eye development. In this assay, populations of homozygous mutant cells are generated next to populations of homozygous wild type cells very early in development. During growth of the eye imaginal disc (the precursor of the adult eye) these clones of cells compete to populate the tissue, resulting in patches of fully mutant (white) and fully wild type (red) tissue in the adult eye (**Figure 4A**). In this assay, *Set8^null^* mutant cells are never present in the adult eye, indicating a strong proliferation defect upon removal of Set8 (**Figure 4B**).

**Figure 4.**
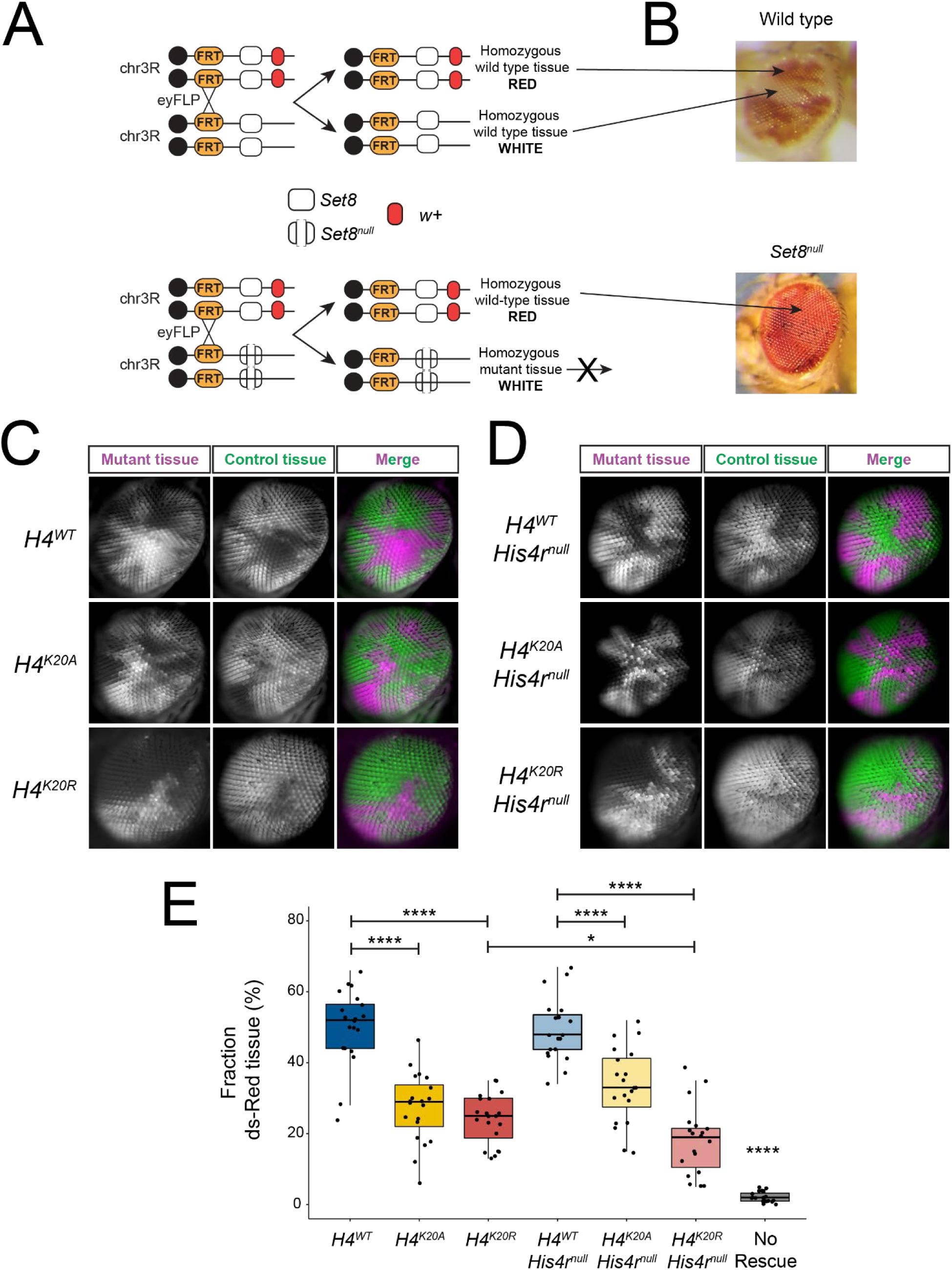
*Set8* but not *H4^K20^* mutant cells fail to proliferate. **A-B)** Diagram of w^+^/w^-^ mosaic eye generation using FLP-FRT mediated mitotic recombination. *Set8* wild type (white box) control experiment (**A**) results in an adult eye with an equal mixture of red and white tissue. *Set8^null^* (open white box) experiment (**B**) results in only red tissue because homozygous *Set8^null^* cells fail to proliferate. **C)** Single optical sections of genetically mosaic eyes composed of clonal populations of cells that are homozygous for *Act5C-GFP* and *HisC*+ (green) or homozygous for *ΔHisC^cadillac^* and lacking *Act5C-GFP* (magenta) rescued by the indicated control or H4K20 mutant transgenes. **D)** Same as (**C**) except in a *His4r^null^* mutant background. Final column shows merged images of the pseudocolored histone mutant clones (magenta) and sister histone wild type clones (green). **E)** Quantification of mosaic eye clones in Figure 4C-D. Boxplot showing the fraction of mutant tissue (magenta) for 19-20 eyes of each genotype (10 males and 9-10 females each). Black dots indicate measurements from individual eyes. Significance was determined by Student’s t-test. **** indicates p < 0.0001, *** indicates p < 0.001, and * indicates p < 0.05.

Assessing the proliferation capacity of histone mutant cells using white versus red marked eye clones was not possible using our previously described histone gene replacement system (McKay et al. 2015), primarily because the *ΔHisC* deletion allele of the endogenous replication-dependent histone genes contains a copy of the *white* gene. To remedy this problem, we engineered a new histone gene deletion (*ΔHisC^cadillac^*) that replaces the *white* gene with *dsRed* driven by the ubiquitously expressed *Act5C* promoter, as well as a wild type *HisC^+^*chromosome marked with an *Act5C-GFP* transgene (Crain, Nevil et al., in preparation). Thus, we can assess proliferation of histone mutant cells (magenta) next to wild type cells (green) during eye development using fluorescence microscopy. We found that *ΔHisC^cadillac^* homozygous mutant cells rescued by one copy of a control *H4^WT^* transgene proliferated normally, resulting in approximately 50% of magenta tissue in the adult eye (**Figure 4C, E**).

In contrast to *Set8^null^* cells, *H4^K20A^* cells were able to proliferate and populate the adult eye but not as well as *H4^WT^*control cells (**Figure 4C, E**). This result is consistent with our previous observation that *H4^K20A^* mutant animals can complete development (Crain et al. 2022). Strikingly, *H4^K20R^* cells proliferate similarly to *H4^K20A^*cells, despite the dramatically different gene expression profiles of these two mutants (**Figure 4C, E**, **Figure 3B, C**). Moreover, a defect in cell proliferation likely does not explain the failure of *H4^K20R^* mutant animals to complete development. Further, *H4^K20A^* or *H4^K20R^* mutant cells lacking the *His4r* gene, and therefore incapable of generating any H4K20me, also generated clones of magenta cells, indicating that the proliferation defect of *Set8^null^*cells is independent of loss of H4K20me (**Figure 4D, E**). Together with our observation that cell proliferation genes are uniquely downregulated in *Set8^null^* animals, we conclude that Set8 functions in cell proliferation independently of H4K20me1 in *Drosophila*, likely via a target other than H4K20.

### Gene expression changes in *l(3)mbt* mutants are not explained by H4K20me1 but correlate with gene expression changes in *Set8^null^* and *H4^K20R^* mutants

Because *Set8^null^* and *H4^K20R^* have disparate developmental phenotypes and gene expression changes that cannot be explained by loss of H4K20me1, we considered other interpretations of our data. Since Set8 and L(3)mbt bind each other directly (Kalakonda et al. 2008) and interact with the H4 tail, mutation of H4K20 might affect recruitment of these proteins to chromatin irrespective of loss of H4K20me. Thus, we asked whether mutation of *l(3)mbt* results in expression changes similar to *Set8^null^* or *H4^K20R^* mutants. We performed RNA-sequencing of +/*Df^ED10966^* (*Df)* hemizygous control, *l(3)mbt^GM76^*/*Df* (hereafter *l(3)mbt^GM76^*), and *l(3)mbt^PBac{Scarless-dsRed}^*/*Df* (hereafter *l(3)mbt^PBac^*) mutant whole third instar larvae at 25°C and at 29°C where the *l(3)mbt* mutant lethality and brain overgrowth phenotypes arise (Bonasio et al. 2010; Janic et al. 2010). *l(3)mbt^GM76^* is a null allele that contains a nonsense mutation in the second of the three L(3)mbt MBT domains (Blanchard et al. 2014; Yohn et al. 2003). *l(3)mbt^PBac^* is a CRISPR-engineered null allele we generated containing a 1.8kb insertion immediately upstream of the *l(3)mbt* start codon.

Loss of L(3)mbt function (*l(3)mbt^GM76^* or *l(3)mbt^PBac^*) significantly disrupts the *Drosophila* transcriptome (**Figure 5A, B**, **Supplemental Table 3**, 1519 up, 358 down; 2938 up, 564 down, respectively). The majority of DEGs are upregulated, consistent with the previously reported transcription repressor function of L(3)mbt (Coux et al. 2018; Kalakonda et al. 2008; Trojer et al. 2007; West et al. 2010). As with *Set8^null^* and *H4^K20R^*, neither the up- nor the down-regulated DEGs in the *l(3)mbt* mutants are enriched for HIGH H4K20me1 genes (**Figure 5A, B**, dark dots, **Supplemental Table 4;** 3.9% in *l(3)mbt^GM76^*, 7.1% in *l(3)mbt^PBac^*). Unexpectedly, we found that removing only one copy of *L(3)mbt* in (+/*Df*) results in many gene expression changes (1716 up, 146 down) relative to true wild type Oregon-R, but with smaller effect sizes than *l(3)mbt^GM76^*or *l(3)mbt^PBac^* (**Figure 5C)**. The *Df* is relatively small, deleting only 28kb including seven genes in addition to *l(3)mbt*: *woc, mrt, TfIIA-L, CG5934, CG5938, CG14260, CG14262.* We found minimal differences in the transcriptomes of *l(3)mbt^GM76^* versus a +/*Df,* suggesting the effect we observed in +/*Df* is due to haploinsufficiency of *l(3)mbt* (data not shown). We only found a small number of gene expression changes in animals raised at 29°C relative to animals raised at 25°C despite a decrease in viability and brain tissue overgrowth at 29°C (Meier et al. 2012), suggesting the phenotypes associated with heat stress are not due to large changes in gene expression.

**Figure 5.**
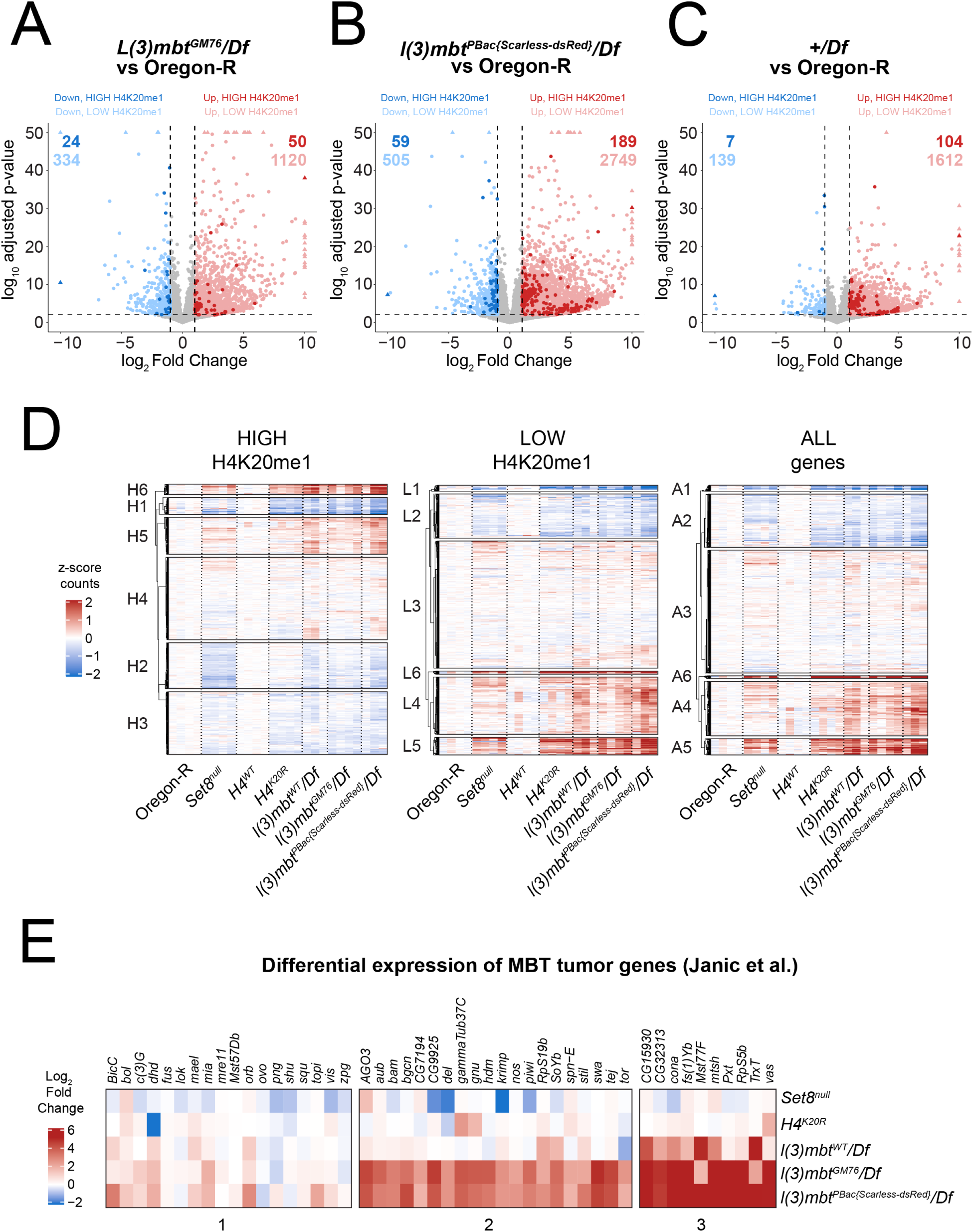
*l(3)mbt, Set8*, and *H4^K20R^* transcriptomes are similar. **A-C)** Volcano plots depicting the relationship between log_2_FC (x-axis) and log_10_ adjusted p-value (y-axis) of gene expression in the indicated comparisons. Blue dots indicate significantly downregulated genes (log_2_FC < −1, FDR < 0.01) and red dots indicate significantly upregulated genes (log_2_FC > 1, FDR < 0.01). Darker shaded dots indicate genes in the HIGH H4K20me1 category. **D)** Clustered heatmaps of average centered normalized counts in the indicated genotypes. **E)** Heatmap of log_2_FC values of in indicated genotypes for genes involved in MBT tumor formation (Janic et al. 2010).

To ask whether loss of L(3)mbt could explain gene expression changes in *Set8^null^* and *H4^K20R^* mutants we compared normalized gene counts in *l(3)mbt* mutants to *Set8^null^*and *H4^K20R^* mutants using k-means clustering. We found several clusters of genes that had similar expression in *l(3)mbt, Set8^null^*and*H4^K20R^*, regardless of H4K20me1 status (**Figure 5D**), suggesting that a subset of gene expression changes in *Set8^null^* or *H4^K20R^* result from altered L(3)mbt function. We observed several clusters that contained genes which were up-regulated in each of the *Set8^null^*, *H4^K20R^*, and *l(3)mbt* mutants, suggesting that expression of these genes share a common mechanism (**Figure 5D**). Janic and colleagues identified a group of 48 genes that were up-regulated in l(3)mbt loss-of-function mutants, resulting in tumors with a characteristic soma-to-germline transition phenotype (Janic et al. 2010). We investigated whether this group of genes were amongst the up-regulated genes in our *l(3)mbt* mutants, and whether *Set8^null^* and *H4^K20R^* shared these gene expression changes. We found that the majority of *l(3)mbt*-tumor genes are upregulated in our *l(3)mbt* mutant datasets (even *+/Df*) but are not significantly changed in *Set8^null^* or *H4^K20R^*mutants (**Figure 5E**). Therefore, disruption of L(3)mbt function from loss of H4K20me1 is likely not substantially contributing to the overgrowth phenotypes associated with MBT tumors.

### L(3)mbt binds the genome independently of H4K20me

Shared gene expression changes in *Set8^null^*, *H4^K20R^*, and *l(3)mbt* mutants might result from loss of L(3)mbt binding to chromatin. The L(3)mbt MBT domains preferentially bind to H4K20me1/2 peptides *in vitro*, (Trojer et al. 2007; Min et al. 2007; Li et al. 2007), although recent *in vivo* studies found that L(3)mbt ChIP-seq peaks colocalize with other methylated histone lysines more than H4K20me1 (Blanchard et al. 2014). We asked whether L(3)mbt recruitment to the genome requires H4K20me using our *H4^K20R^, His4r^null^* genotype lacking all H4K20. Since no L(3)mbt antibodies were available to us, we engineered N-terminal GFP- and FLAG-tagged alleles at the endogenous *l(3)mbt* locus to assess genome-wide binding of L(3)mbt (**Figure 6A**). Animals expressing only GFP- or FLAG-tagged L(3)mbt are viable and display no obvious phenotypic abnormalities. Using anti-GFP- or anti-FLAG antibodies we found that our epitope tagged L(3)mbt proteins are expressed in 3^rd^ instar larval brains, adult ovaries, wing imaginal discs in patterns and levels consistent with previous studies (**Figure 6A**) (Coux et al. 2018; Meier et al. 2012; Richter et al. 2011; Yamamoto-Matsuda et al. 2022).

**Figure 6.**
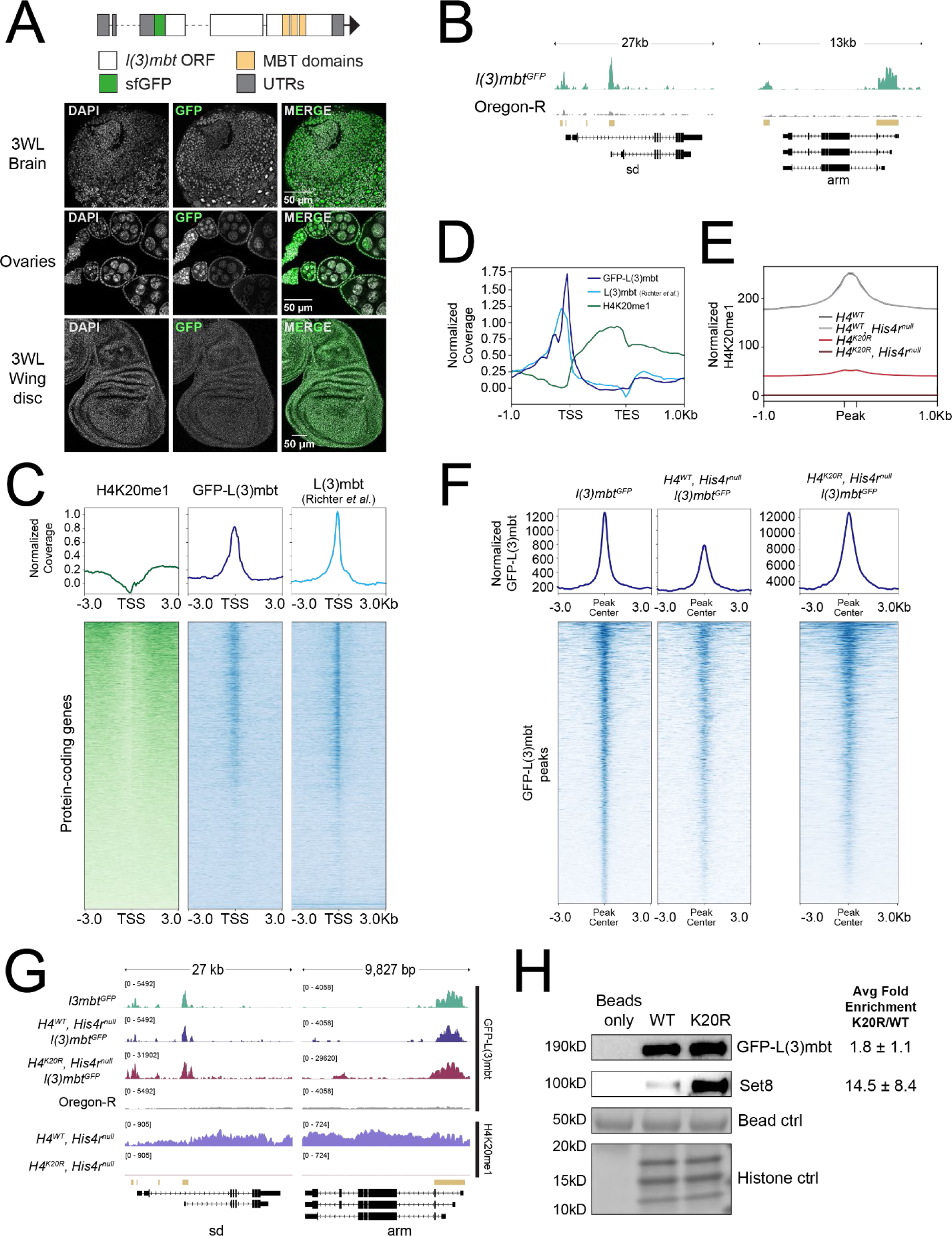
L(3)mbt binds the genome independently of H4K20me. **A)** Diagram of GFP-tagged *l(3)mbt* allele generated using CRISPR (top). Confocal images of GFP-L(3)mbt accumulation in the nuclei of third instar larval brains, adult female ovaries, and third instar larval wing discs. **B)** Representative locus depicting normalized GFP-L(3)mbt CUT&RUN coverage. Gold bars indicate peaks of GFP signal in *l(3)mbt^GFP^* (top) relative to Oregon-R control (bottom) as defined by merged 150bp sliding windows. **C)** Heat (bottom) and summary (top) metaplots of z-normalized wing disc H4K20me1 CUT&RUN, wing disc GFP-L(3)mbt CUT&RUN, and L(3)mbt ChIP-seq (Richter et al. 2011) signal at protein-coding genes. Plots are centered at the TSS and flanked by 3kb of unscaled sequence. Each row represents a single gene, and genes are ordered by mean H4K20me1. Summary plot depicts mean signal for H4K20me1 (green), GFP-L(3)mbt (blue), or L(3)mbt (light blue) at each position in the metaplot (50bp bins). **D)** Summary metaplot of z-normalized H4K20me1 (green), GFP-L(3)mbt CUT&RUN (blue) and L(3)mbt ChIP-seq (light blue) at HIGH H4K20me1 genes. **E)** Summary metaplot of spike-in normalized H4K20me1 CUT&RUN coverage at H4K20me1 peaks in the indicated genotypes. Each peak is scaled to 200bp and flanked by 1kb of unscaled sequence. Summary plot depicts mean signal at each position in the metaplot (50bp bins) for all peaks. **F)** Summary (top) and heat (bottom) metaplots of spike-in normalized GFP-L(3)mbt CUT&RUN signal. Plots are centered at the GFP-L(3)mbt peak, and each peak is flanked by 3kb of unscaled sequence. Each row represents a single peak, and peaks are ordered by mean GFP-L(3)mbt in *l(3)mbt^GFP^***G)** Representative locus depicting spike-in normalized GFP-L(3)mbt and H4K20me1 CUT&RUN signal. Gold bars indicate peaks of GFP signal in *l(3)mbt^GFP^* relative to Oregon-R control as defined by merged 150bp sliding windows. **F)** Western blot using anti-GFP (top) or anti-Set8 (middle) antibodies following recombinant nucleosome binding assay with nuclear lysate from third instar larvae and either wild type (WT) or H4K20R (K20R) recombinant nucleosomes. Histone proteins from Swift protein stain are used as loading control. Average binding of Set8 and GFP-L(3)mbt on K20R vs. WT nucleosomes ± standard deviation from 3 biological replicates shown on right.

We performed CUT&RUN in third instar wing imaginal discs that express only GFP-L(3)mbt from the endogenous locus. We identified regions of the genome bound by L(3)mbt by measuring enrichment of GFP-L(3)mbt signal over an Oregon-R control (which lacks GFP). We found that L(3)mbt accumulates at 4,052 well-defined peaks across the genome that are preferentially located at transcription start sites of genes enriched with H4K20me1 (**Figure 6B, C, D**). Despite the correlation with H4K20me1-enriched genes, L(3)mbt is not enriched in gene bodies, unlike H4K20me1 (**Figure 6D**). To corroborate this observation, we also analyzed the location of peaks in a previously published L(3)mbt ChIP-seq dataset from third instar larval brains (Richter et al., 2011) and found that L(3)mbt is enriched at transcription start sites of HIGH H4K20me1 genes in both datasets (**Figure 6C, D)**.

We next performed anti-GFP-L(3)mbt CUT&RUN in a genotype that lacked all modifiable H4K20 (*H4^K20R^, His4r^null^*). We found that *H4^K20R^, His4r^null^* third instar wing discs have a genome-wide ablation of H4K20me1 while removal of *His4r* alone (*H4^WT^, His4r^null^*) does not detectably affect total H4K20me1 levels (**Figure 6E**). Since H4K20me increases the affinity of L(3)mbt for H4 tail peptides in vitro (Trojer et al. 2007; Li et al. 2007; Min et al. 2007), we hypothesized that loss of modifiable H4K20 would result in a reduction of GFP-L(3)mbt signal at GFP-L(3)mbt peaks. We observed that GFP-L(3)mbt is recruited to the genome at most of its binding sites when all H4K20 is mutated to arginine (**Figure 6F, G**). Unexpectedly, we also observed a >10-fold increase of GFP-L(3)mbt binding across all peaks in *H4^K20R^, His4r^null^* compared to *H4^WT^, His4r^null^* (**Figure 6F, G**). These data indicate that mutation of H4K20 to Arg results in altered, rather than the prevention of, accumulation of GFP-L(3)mbt on chromatin.

To determine how GFP-L(3)mbt directly associates with H4K20R nucleosomes, we performed in vitro binding assays using H4K20R recombinant nucleosomes (Skrajna et al. 2020). We found that GFP-L(3)mbt exhibited a modest increase (1.8 ± 1.1) in binding to H4K20R nucleosomes compared to WT nucleosomes (Figure 6H). Interestingly, we also observed a 14.5-fold (± 8.4) increase in the binding of Set8 to H4K20R nucleosomes relative to WT nucleosomes (Figure 6H). These data suggest that the Lys to Arg substitution results in neomorphic effects likely due to increased H4 tail binding of L(3)mbt, Set8, and possibly other proteins, a phenomenon also found in onco-histone mutations such as H3K27M (Sahu and Lu 2022). Together these data indicate that H4K20me is not necessary for L(3)mbt to bind chromatin in vivo.

## Discussion

By combining genetic and genomic approaches in *Drosophila melanogaster* we provide evidence that H4K20me1 is dispensable for key functions of Set8, including regulation of gene expression and cell proliferation, that have been previously attributed to H4K20me1. We also demonstrate that L(3)mbt functions in gene expression independently of binding H4K20me.

### Set8 and L(3)mbt function in gene expression independent of H4K20me1

Our data do not support a model whereby the primary functions of Set8 and L(3)mbt are mediated through the deposition and recognition of H4K20me1, respectively. We found that *Set8* and *l(3)mbt* null mutations have a greater effect on the *Drosophila* transcriptome than *Set8^RG^*, *H4^K20A^*, or *H4^K20R^*mutants. Remarkably, *Set8^RG^* and *H4^K20A^* mutants have minimal gene expression changes – none of which are in genes with the highest coverage of H4K20me1 – despite a strong genome-wide reduction in H4K20me1. More genes change in expression in *H4^K20R^* mutants than in *H4^K20A^* mutants, but many of these changes do not correlate with gene expression changes in *Set8^null^*. Moreover, we demonstrate that cells lacking all modifiable H4K20 (*H4^K20A^, His4r^null^* or *H4^K20R^, His4r^null^*) can proliferate in stark contrast to *Set8^null^* cells which cannot, indicating that Set8, not its catalytic activity on H4K20, is required for cell proliferation.

Similarly, we found that H4K20me is dispensable for L(3)mbt recruitment to the genome, including promoter-associated peaks. Methylated lysine residues other than H4K20me could help recruit L(3)mbt to specific promoters, consistent with recent work reporting L(3)mbt co-enrichment with several different methylated histone residues at promoters (Blanchard et al. 2014) A similar situation occurs with other MBT domain-containing proteins such as Sfmbt (Klymenko et al. 2006). Instead, H4K20me could be required for positioning of L(3)mbt on chromatin or chromatin compaction after being recruited via other mechanisms (Min et al. 2007; Trojer et al. 2007; Li et al. 2007; Blanchard et al. 2014). We conclude that many, if not most, of the critical cellular and developmental functions of Set8 and L(3)mbt in *Drosophila* are likely mediated through targets other than H4K20.

We were surprised to find that H4K20me1 had little impact on gene expression, given that H4K20me1 is enriched in the bodies of expressed genes in both flies (this study, (Lv et al. 2016)) and humans (Barski et al. 2007; Beck et al. 2012; Congdon et al. 2010). Indeed, H4K20me1 is among the histone PTMs most highly correlated with active transcription (Wang et al. 2008). Set8 interacts with elongating RNA Polymerase II (Li et al. 2011), so H4K20me1 accumulation in gene bodies may be a consequence of the Set8/RNA Pol II interaction without playing a regulatory role in transcription. Although we don’t observe gene expression changes associated with mutation of H4K20 in our whole larvae datasets, our analysis could lack the power to detect H4K20me1-dependent tissue-specific gene expression changes. For instance, H4K20me1 covers genes that are uniquely expressed in specific cell lines (Beck et al. 2012). H4K20me1 might promote a permissive environment in which tissue-specific transcription factors function more efficiently or help counteract other repressive chromatin domains. For instance, Lv et al. observed that Pc+, H3K27me3+, H4K20me1+ genes were expressed whereas Pc+, H3K27me3+, H4K20me1- were repressed (Lv et al. 2016). Another proposed role of H4K20me1 is recruitment of the MSL complex to the TSS to release paused polymerase into productive elongation (Kapoor-Vazirani and Vertino 2014; Nikolaou et al. 2017). Since all our data were collected from female larvae, we were not able to address this mechanism. Nevertheless, our data demonstrate that H4K20me has no uniform genome-wide regulatory role, either positively or negatively, in gene expression in *Drosophila*.

### Mechanisms of H4K20 methylation

We detected low levels of H4K20me1 in *Set8^null^* animals by CUT&RUN, which was unexpected as Set8 is currently the only known H4K20 mono-methyltransferase, and previously we did not detect H4K20me1 signal by western blot in *Set8^null^* mutants (Crain et al. 2022). We detected no H4K20me1 CUT&RUN signal in *H4^K20R^*, *His4r^null^* animals, which contain no modifiable H4K20, indicating that the H4K20me1 antibody does not recognize other histone PTMs. Although our data cannot exclude that the H4K20me1 antibody binds also to unmodified H4K20, they raise the possibility of another H4K20 mono-methyltransferase in flies. Karachentsev et al. also reported residual H4K20me1 in *Set8^null^* salivary glands by immunofluorescence that they hypothesized was due to another H4K20 mono-methyltransferase or stabilization of the mono-methyl mark over multiple cell divisions (Karachentsev et al. 2005). One possibility is that the H4K20 di- and tri- methyltransferase Suv4-20 can generate H4K20me1 in the absence of Set8. Three studies reported that Suv4-20 preferentially utilizes H4K20me1 as a substrate *in vitro* but can also produce H4K20me1 (Weirich et al. 2016; Yang et al. 2008; Southall et al. 2014). Another possibility is the production of H4K20me1 by demethlyation of long-lived pools of H4K20me2,3 that are derived from maternally deposited Set8 in *Set8^null^* mutant animals. Although not yet identified in flies, H4K20 demethylase enzymes are present in other metazoans, including PHF8 in zebrafish and mammals and DPY-21 in C. elegans (Qi et al. 2010; Brejc et al. 2017; Gu et al. 2016a). Regardless of the source of this small amount of H4K20me1 in *Set8^null^* mutants, our results clearly indicate that Set8 is responsible for the bulk of H4K20me1 in *Drosophila*.

### The *H4^K20A^* and *H4^K20R^* mutant phenotypes differ

H4K20me1 has been implicated in many essential nuclear processes, and thus our previous observation that animals expressing only unmodifiable H4K20A histones can complete development was surprising (Crain et al. 2022). This result is consistent with the small number of gene expression changes in H4K20A mutants we report here. However, only 20% of *H4^K20A^*animals, and no *H4^K20R^* animals, reach adulthood, indicating that H4K20 is important for development (Crain et al. 2022). The more extensive gene expression changes *H4^K20R^* animals could explain why they die during development. These changes do not overlap substantially with gene expression changes in *Set8^null^*mutants. By contrast, we observed similar gene expression profiles in *H4^K20R^*and *l(3)mbt* mutants. These changes are not driven by loss of H4K20me but rather by mutating histone H4K20 to Arg. Thus, these data indicate that H4K20me is not essential for *Drosophila* development, but do not explain the phenotypic differences between *H4^K20A^* and *H4^K20R^*mutants or which processes are affected by mutating H4K20.

We suspect that the Lys to Ala and Lys to Arg amino acid changes in the H4 tail disrupt or change the interaction of nucleosome binding proteins irrespective of the absence of H4K20me. These proteins include Set8 and L(3)mbt, as we observed binding of L(3)mbt to both the H4K20R mutant genome and purified H4K20R nucleosomes, and increased binding of Set8 to purified H4K20R nucleosomes. The H4K20 mutations may also affect te H4 tail conformation and thus the availability *in vivo* of the H4 tail to chromatin binding proteins. Regardless, alteration of Set8 or L(3)mbt H4 tail binding alone is insufficient to explain the phenotypes of *Set8^null^* and *l(3)mbt* mutant animals because they are so different than the *H4^K20^* mutant phenotypes (Crain et al. 2022). Future determination of the H4 tail and/or broader nucleosome interactome in *H4^K20A^* and *H4^K20R^*mutants should be informative in this regard.

Our study illustrates the power of employing genomic analyses in a genetically tractable organism like *Drosophila melanogaster* to deconvolve the complex relationship between a writer (Set8) and a reader (L(3)mbt) of a particular histone PTM (H4K20me). Notably, as the chromatin field continues to identify pleiotropic functions of writer and reader proteins, revisiting other histone PTM/writer/reader paradigms using a similar type of analysis may prove fruitful in untangling the functions of histone PTMs in various biological processes.

## Materials and Methods

### GFP-L(3)mbt Immunofluorescence

Cuticles from *l(3)mbt^GFP^* larvae were inverted and fixed in 3.7% paraformaldehyde for 25 min, washed, then blocked in 500μL 5% NGS in PBS for 30 min prior to staining with Abcam α-GFP Rb ab6556 1° (1:1000) ON at 4°C followed by Alexa rabbit 488 2° (1:1000) at RT for 2h. Tissues were dissected off cuticles, mounted on a glass slide with a glass coverslip in 11μL Prolong and left in the dark overnight before imaging on a Leica SP8 confocal microscope. Ovaries were dissected from 3-day-old adult females then fixed and stained as above.

### Larvae collection and RNA extraction

Four replicates of 8 third instar wandering larvae of each genotype were homogenized in TRIzol (Invitrogen) and flash frozen in liquid nitrogen. RNA was isolated using the Direct-zol RNA miniprep kit (Zymo).

### RNA-seq library preparation and sequencing

RNA-seq libraries were prepared using the Universal Plus Total RNA-seq with NuQuant (Tecan). Number of cycles required for library amplification was determined empirically using qPCR. DNA concentrations, fragment size distributions, and quality were determined using Qubit and 4150 Aglient Tapestation. Libraries were pooled and sequenced PE100 on a Nova-seq.

### RNA-seq

#### Differential expression and RNA abundance

Paired-end FASTQ files from 3-4 replicates of each genotype were passed to the quant function within Salmon (Patro et al. 2017) in mapping-based mode with the parameters *validateMappings, seqBias, eVBOpt, numBootstraps 30* and transcript indexes and decoys generated from the dm6 *Drosophila* genome build. Quant files were imported into R using Txiimport (Soneson et al. 2015). Genotypes for all replicates were confirmed using sequencing data. Replicates with reads that did not match the intended genotype were discarded and not used for further analyses. For RNA abundance, transcripts per million (TPM) counts were obtained from Salmon output. Normalized counts from all replicates of a given genotype were averaged and plotted using ggpubr R package with the parameter outlier.shape = NA. Differential analysis was performed using DESeq2 (Love et al. 2014). Log_2_FC values were shrunk using the ashr method (Stephens 2017). Heatmaps were generated using ComplexHeatmap (Gu et al. 2016b). Boxplots generated using ggplot (Wickham 2016).

### CUT&RUN

20 third instar larval wing discs per replicate were dissected and processed as previously described (Uyehara et al. 2022) with rabbit α-GFP (1:100, Rockland 600-401-215) or mouse α-H4K20me1 (1:100, ThermoFisher MA5-18067) and pAG-MNase (1:100, UNC core, (Salzler et al. 2023)).

### Library Preparation and Sequencing

DNA libraries were prepared using ThruPLEX® DNA-Seq Kit (Takara) and DNA Unique Dual Index Kit with associated protocols. DNA concentration, fragment distribution, and quality were determined by Qubit and 4150 Agilent TapeStation. Libraries were pooled and sequenced PE75 on a NextSeqP3 or NextSeq2000.

### CUT&RUN Sequencing Data Analysis

#### Data processing

Replicates from each genotype were processed using a Snakemake pipeline (https://github.com/snystrom/cutNrun-pipeline.git). Adapters were trimmed with bbduk, reads were aligned to dm6 with bowtie2 (Langmead and Salzberg 2012), converted to Bam format and reads with a quality score less than 30 for GFP CUT&RUN and 5 for H4K20me1 CUT&RUN were removed via samtools (Li et al. 2009), and duplicate reads were kept for GFP CUT&RUN and removed for H4K20me1 CUT&RUN. Bam files were generated, sorted, and converted to bed format with bedtools (Quinlan and Hall 2010)

#### Peak calling

The following was performed using the csaw package (Lun and Smyth 2016) unless stated otherwise. Reads from Oregon-R (H4K20me1) and Oregon-R (no primary) were binned into 150bp windows with a 50bp slide. Background was calculated by binning reads into large 10kbp windows and 150bp windows were retained only if they were log_2_(2) higher than background. Window counts for each replicate were normalized for compositional bias and differential binding was assessed using edgeR (Robinson et al. 2010). Windows that were significantly enriched (log_2_FC > 1 and FDR < 0.05) in Oregon-R (H4K20me1) over Oregon-R (no primary antibody) were merged if they within 1kb of each other and FDR < 0.05. Subsequent merged windows were saved to bed format and used for downstream analyses. Peaks in GFP-L(3)mbt over Oregon-R negative control were determined as in H4K20me1 with the following exceptions: 1) log_2_FC > 3 significance cutoff 2) Peaks within 250bp were merged.

#### Average signal plots

For figure 1D: Coverage files for 3 biological replicates of H4K20me1 Oregon-R wing disc and no primary Oregon-R CUT&RUN and 2 biological replicates of H4K20me1 Oregon-R whole larvae ChIP-seq and input controls ((modENCODE Consortium et al. 2010), GSE47254) were normalized by reads per genome content (RPGC) and averaged using deeptools BigWigAverage (Ramírez et al. 2016). The ratio of each averaged H4K20me1 file over control was calculated using deeptools BigWigCompare (Ramírez et al. 2016) and then z-normalized.

For Figures 2A and 6E: Individual genome coverage files (three replicates; two for *H4^WT^*, *H4^K20A^*, and *H4^K20R^, His4r^null^*; see below) for each genotype were normalized using yeast spike-in controls by the Spike-in normalized Reads Per Million mapped reads in the negative Control (SRPMC) method (DeBerardine et al. 2023). The ratio of fly to yeast reads was calculated (reads per spike-in, RPS), RPS values between each sample and its control was calculated (Relative signal), and the relative signal for each sample was RPGC scaled. The SRPMC scaling factor was used in bedtools genomeCoverage (Quinlan and Hall 2010) to produce scaled bedgraph files which were subsequently converted to BigWig coverage files using ucsctools wigToBigWig. *H4^WT^* rep3 was removed due to an exceptionally high yeast spike in count, *H4^K20A^* rep1 was removed because it contained wild type reads at H4K20, and *H4^K20R^, His4r^null^* rep2 was removed due to exceptionally low read count. The remaining replicates of each genotype were averaged using deeptools bigWigAverage (Ramírez et al. 2016) using 1bp bins. Metaplots were generated using computeMatrix and plotHeatmap functions in deeptools (Ramírez et al. 2016). Two biological replicates of each *l(3)mbt* genotype for α-GFP CUT&RUN were processed as above.

### Mitotic eye clone generation and quantification

*eyFLP; Actin5C-GFP/CyO;* + or *eyFLP; Actin5C-GFP/CyO; His4r^null^* females were crossed to *yw; ΔHisC^cadillac^/CyO; 12xHTG* or *yw; ΔHisC^cadillac^/CyO; 12xHTG*, *His4r^null^* males. 19-20 adults (9-10 males and females each) were aged 1-2 days after eclosion, placed in a 96-well dish containing molten 1% agarose, and cooled to solidify. Images were obtained on a Leica M205 FCA fluorescent microscope using GFP and RFP band pass filters. Quantification of mutant clone size was determined using FIJI. Area of the eye from stacked RGB eye images was manually cropped to an ellipse using the selection tool. Eye clones were then white pseudocolored using Color Histogram. The image was then converted into a binary image, followed by restoration of the eye area selection. Finally, the area of the eye covered by mutant clones within the eye area selection was measured and recorded.

### Recombinant nucleosome binding assays

Nuclei from 100-120 *l(3)mbt^GFP^* third instar wandering larvae were collected as previously described (Leatham-Jensen et al. 2019) Nuclei pellets were resuspended in 500µL BB420 buffer (20mM HEPES pH 7.5, 420mM NaCl, 1mM DTT, 0.1mM EDTA, 10% glycerol, 0.1% NP-40), homogenized 10x with a Dounce homogenizer, then centrifuged at 17k x g for 30 min. Supernatant was moved to a new 1.5mL tube and brought to a final salt concentration of 120mM (BB120). The H4K20R mutant was cloned using site directed mutagenesis. Nucleosomes containing FLAG-tagged H2A and either WT H4 or H4K20R were assembled, and binding assays were performed as previously described (Skrajna et al. 2020). Briefly, 20µL of anti-FLAG-conjugated magnetic beads (Millipore Sigma) were incubated with 20µg of FLAG-tagged WT or H4K20R nucleosomes or equivalent volume of BB120 (beads only control) for 2 h at 4°C. Beads were washed 3x with BB120. Nuclear lysate was split equally between the three tubes and the volume was increased to 500uL with BB120 prior to incubation for 2h at 4°C. Beads were washed 2x with BB120, followed by a 30min wash in BB120 at 4°C. Following the last wash, bound proteins were eluted with 25µL gel loading buffer and samples were boiled for 5min. Samples were run on a 4-20% gradient SDS-PAGE gel (Bio-Rad) and transferred to a nitrocellulose membrane (Bio-Rad Quick transfer). Total protein staining was performed using Swift protein stain (G-Biosciences). Membranes were blocked in 5% milk in TBS-Tween for 1h prior to incubation with α-GFP (1:1000, Rockland) or α-Set8 (1:1000, Novus Biologicals) primary antibody overnight at 4°C, followed by rabbit HRP secondary (1:10,000) at room temperature for 1h. Blots were incubated in SuperSignal West Pico Chemiluminescent Substrate (Fisher) for 5min then imaged on an Amersham imager (GE). Entire blot was autoscaled for brightness/contrast prior to quantification by densitometry in FIJI.

## Acknowledgements

We thank Markus Nevil and Harmony Salzler for critical reading of the manuscript, Gabrielle Budziszewski for production of the H4K20R histone plasmid, Alexsandra Skrajna for providing reagents, protocols, and troubleshooting advice for nucleosome binding assays, and the UNC HTSF sequencing core for RNA-seq. This work was supported by NIH T32GM007092 to ATC and NIH grants R35GM133498 to RKM and R01GM124201 and R35GM145258 to RJD.

## Author contributions

ATC and RJD conceptualized the study. ATC and MBB performed experiments and data analysis. CAH contributed to CUT&RUN and RNA-seq experiments. ATC and MBB performed bioinformatic analyses. RKM provided expertise, troubleshooting advice, and reagents for nucleosome binding assays. MH assembled recombinant nucleosomes, including extensive troubleshooting. ATC and RJD wrote the manuscript.

